# Substrate properties and actin polymerization speed dictate universal modes of cell migration: gripping, slipping, and stick-slip

**DOI:** 10.1101/2025.06.20.660727

**Authors:** Yiyang Ye, Jie Lin

## Abstract

Understanding how cells sense mechanical cues and regulate migration is crucial in the development, fibrosis, and oncogenesis processes. However, a comprehensive physical picture of cell migration remains lacking, given the diverse environmental properties and cell physiologies. Here, we generalize the motor-clutch model to the whole-cell level and systematically investigate the effects of substrate stiffness, friction, and actin polymerization speed on cell migration. We unveil three distinct migration modes: gripping, slipping, and stick-slip. Notably, stiffness sensing occurs exclusively in the stick-slip mode, which requires a low substrate stiffness and a minimum actin polymerization speed as necessary conditions. Intriguingly, the optimal substrate stiffness that maximizes the migration speed is inversely proportional to the actin polymerization speed. Moreover, the maximal speed only depends on the nature of the clutch molecules, independent of substrate properties. We reveal the boundary criteria between the three migration modes and demonstrate that fast- and slow-migrating cells can coexist in an isogenic cell population without the need for biochemical feedback loops.

---

Cellular behaviors such as migration [1–3], spreading [4], and differentiation [5, 6] are sensitive to cell-matrix interactions and crucial for developmental processes, tumorigenesis [7] and wound healing [8]. During cell migration, force transmission begins with adaptor proteins linked to actin and proceeds through integrins, which bind to ligands on the extracellular matrix (ECM) [9, 10]. Polymerization of filamentous actin (F-actin) at the leading edge leads to retrograde actin flow, which can be converted to cell migration relative to the substrate through focal adhesions [11–13]. In the meantime, myosin II localized at the cell rear disassembles the actin network together with ADF/cofilins, which allows the recycling of actin monomers [14, 15].

Multiple mathematical models have been proposed to understand cell migration from different perspectives [16]. Among them, the motor-clutch model is one of the most illuminating ones [17–25]. The model was first proposed to explain how substrate stiffness affects filopodia traction dynamics in which molecular clutches link F-actin to the substrate and mechanically resist the retrograde flow [26]. As our understanding of mechanosensing improves, the motor-clutch model has been extended to study the effects of integrin subtypes [27], substrate stiffness gradients [28], ECM viscoelasticity [29–33], adhesion growth [34], spatial sensing of ligands [35], effects of force loading rate [36], etc. Despite the vast applications of the motor-clutch model in various experimental conditions, several key issues remain.

First, cellular physiologies, e.g., the actin polymerization speed, can vary significantly across different cell types; indeed, migration speed varies from less than 1 *µ*m/min for fibroblasts to 15 *µ*m/min for keratocytes [37]. Second, substrate properties, e.g., the stiffness, can vary by orders of magnitude across different experiments [28]. Third, even for an isogenic population in the same environment, experimental observations of freely migrating cells have revealed significant cell-to-cell variability, including a slow-moving subpopulation [38, 39]. However, most theoretical studies of the motor-clutch model are computational and/or involve a dozen variables, in which parameter values are chosen for a particular cell type and experimental setup; therefore, the universality and generalizability of their conclusions are unclear [23]. Thus, an analytically treatable model connecting cell migration to cellular physiologies (e.g., actin polymerization speed) and environmental properties (e.g., substrate stiffness) and capturing phenotypic variability is fundamentally essential, which is presented in this work.

Here, we study the motor-clutch model at the whole-cell level, which allows us to explicitly relate the migration speed to the actin polymerization speed, the substrate stiffness, and the substrate friction. Using a mean-field approximation, we significantly simplify the model to a system with two degrees of freedom: the engagement probability of clutches (*P*_*b*_) and the total traction force (*F*). We obtain the phase diagram of cell migration and reveal three distinct migration modes: gripping, slipping, and stick-slip. The transition mechanisms between different modes are rich, including saddle-node bifurcation, discontinuous and continuous Hopf bifurcation [40].

We highlight several key insights gained from our model: (1) stiffness sensing only occurs in the stick-slip mode of cell migration where the optimal substrate stiffness maximizing migration speed is inversely proportional to the actin polymerization speed, but the maximal migration speed only depends on the properties of molecular clutches; (2) the migration speed and traction force show a biphasic dependence on the actin polymerization speed; (3) the period of the stick-slip mode is proportional to the clutch number, the inverse of substrate stiffness, and the inverse of actin polymerization speed; (4) different migration modes, e.g., fast and slow-moving cells, can coexist in an isogenic population without relying on biochemical feedback loops.

An interesting implication of our model is that the contraction force exerted by Myosin II on the actin filament is irrelevant to the migration speed of a motile cell, as it is an internal force within the cell, and the only external force acting on the cell is from the substrate. Our model rationalizes previous puzzling observations that myosin activity does not affect migration [15, 41–43], and is consistent with the observations that cell migration could be powered solely by actin assembly [44].

## Model description

We introduce a modified motor-clutch model to study a migrating cell moving in a fixed direction, which we denote as the cell migration model in the following (Figure 1). F-actin polymerizes at the leading edge with a constant speed *v*_*p*_. Meanwhile, the speed of the retro-grade actin flow relative to the substrate is *v*_*f*_, and the engagement of molecular clutches to the F-actin resists the retrograde flow [45]. The engagement of clutch *i* with F-actin is characterized by a constant association rate *r*_*on*_ and an unbinding rate, which increases exponentially with force *F*_*i*_ exerted on clutch *i*: 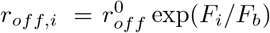, where *F*_*b*_ is a characteristic force for clutch breakage [46].

**FIG. 1.**
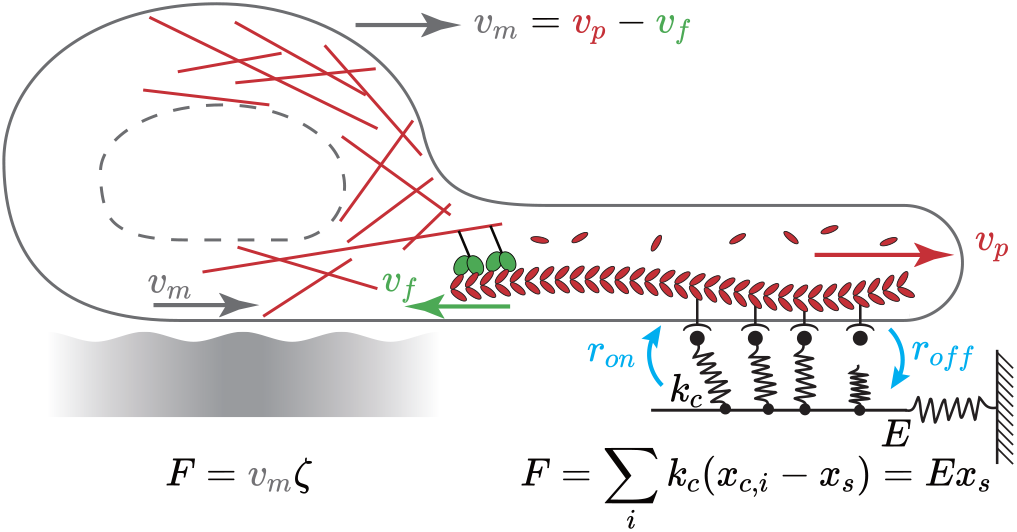
A schematic of the cell migration model. We consider a cell migrating towards the right side with migration speed *v*_*m*_, related to the actin polymerization speed *v*_*p*_ and the retrograde speed of F-actin *v*_*f*_ . The clutches can engage with F-actin and resist its retrograde flow, and they are modeled as springs with spring constant *k*_*c*_. The substrate is modeled as a spring with a spring constant *E*. The total traction force is the sum of the forces on all clutches, which also sets the substrate displacement *x*_*s*_. Furthermore, the total traction force is also proportional to the migration speed as *F* = *v*_*m*_*ζ* where *ζ* is the friction coefficient of the substrate.

The force on each engaged clutch is proportional to its elongation, *F*_*i*_ = *k*_*c*_(*x*_*c*,*i*_ − *x*_*s*_) where *k*_*c*_ is the spring constant of the clutch, *x*_*c*,*i*_ is the displacement of the clutch’s end bound to the F-actin. The substrate displacement, denoted by *x*_*s*_, is set by the total traction force exerted by the migrating cell: *F* = *Ex*_*s*_ = ∑_*i*_ *F*_*i*_, where *F*_*i*_ = 0 for unengaged clutches. The spring constant *k*_*c*_ should be regarded as the overall spring constant of the series of the ligand, integrin, and other proteins involved in the force transmission through a single clutch [47].

By exerting backward traction force on the substrate, the cell migrates forward with a migration speed *v*_*m*_ = *F/ζ*, where *ζ* is the friction coefficient between the cell and substrate. Our model is consistent with the observations that traction force is localized at the cell front while the back of the cell moves passively against the substrate, generating the friction force (Figure 1) [44]. The retrograde flow speed *v*_*f*_ is related to the migration speed *v*_*m*_ and the actin polymerization speed *v*_*p*_ through *v*_*f*_ = *v*_*p*_ − *v*_*m*_. If the cell moves without relative displacement between the F-actin and substrate, *v*_*f*_ = 0 and *v*_*m*_ = *v*_*p*_, it moves at its highest possible speed [15]. We provide an estimation of the typical values of the parameters in Table S1 of the Supplemental Material.

### Mean-field model

We introduce *P*_*b*_ as the fraction of engaged clutches, *P*_*b*_ = ∑_*i*_ *s*_*i*_*/N*, where *N* is the total number of clutches, *s*_*i*_ = 1 for engaged clutches and *s*_*i*_ = 0 for unengaged ones. The time derivative of *P*_*b*_ is 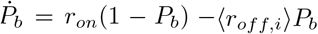 where 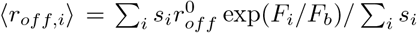. In the mean-field model, the force fluctuation among clutches is neglected, and ⟨*r*_*off*,*is*_⟩ is replaced by 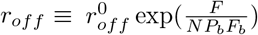. Using the same approximation, the mean-field equation of the total traction force becomes 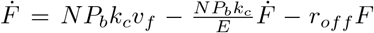, where we have used the fact that a newly engaged clutch bears zero force, i.e., *x*_*c*,*i*_ = *x*_*s*_ right after engagement (see Section A of Supplemental Material for a detailed derivation from the full stochastic model). In the following, we analyze the dimensionless equations of the mean-field model:

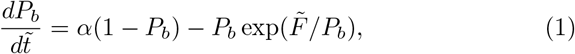

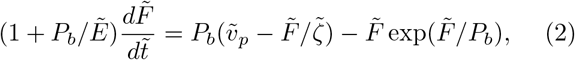

Here, 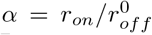. We take 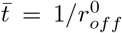 as the time unit, 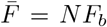 as the force unit, and 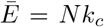 as the stiffness unit. Therefore, the dimensionless actin polymerization speed becomes 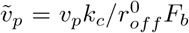 and the dimensionless friction coefficient becomes 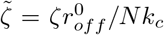. The dimensionless equations unify the behaviors of different cell types and experimental conditions.

We set the time derivatives of Eqs. (1, 2) as zero to analyze the steady-state properties and obtain

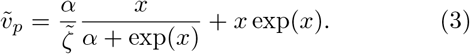

Here, 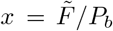 defines the averaged force on a single clutch, with the steady-state binding probability given by *P*_*b*_ = *α/*[*α* + exp(*x*)]. In steady states, Eq. (3) means that 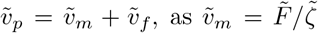 corresponding to the first term on the right hand side; thus, the retrograde flow speed is 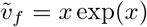. We highlight two key features of Eq. (3). First, a critical substrate friction 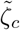 exists below which the left side of Eq. (3) is non-monotonic, and three solutions may exist in which the middle solution must be unstable (Figure 2B, Supplemental Material Section B and Figure S1A). Second, the solutions of Eq. (3) are independent of the substrate stiffness 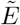, but 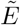 influences the stability of these solutions, as we show later.

**FIG. 2.**
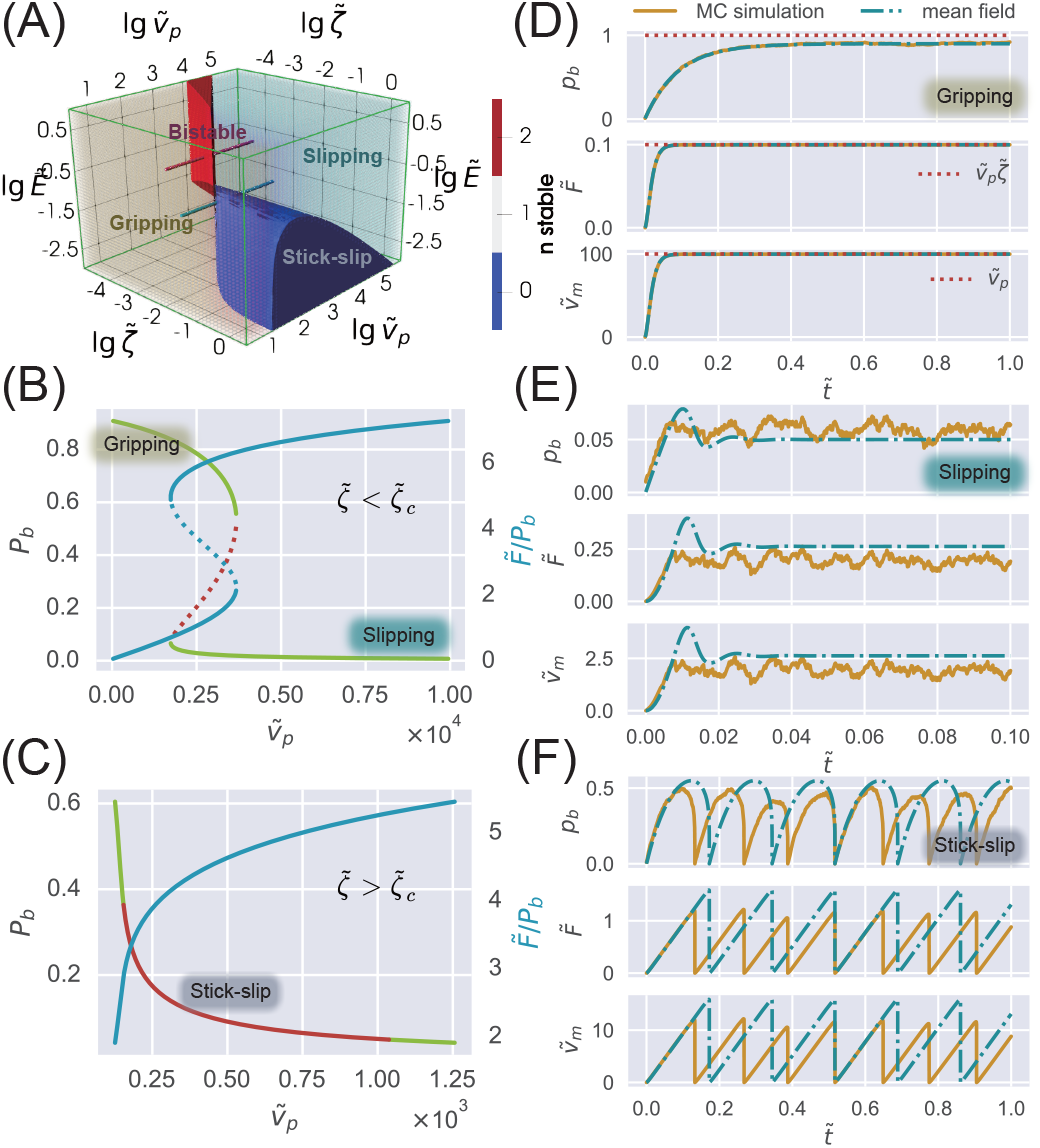
Modes of cell migration. (A) Phase diagram of migration modes as a function of 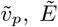 and 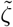. The red area denotes the bistable phase, where slipping and gripping modes coexist. The solid blue area identifies the stick-slip mode. The remaining monostable region comprises two stable modes: the gripping mode (yellow) and the slipping mode (light blue), with a continuous transition between them. (B) The fraction of engaged clutches *P*_*b*_ (green line) and the force per clutch 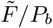 (blue line) vs. 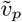 according to Eq. (3) under the condition 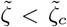 such that three solutions may exist at most. The change of 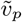 can be considered as moving along the red rod in (A). The dotted lines represent the unstable middle solution. (C) The same as (B) but for 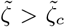 in which only one solution exists for Eq. (3), which is unstable in the red regime. The change of 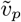 can be considered as moving along the blue rod in (A). (D-F) A comparison of direct simulations of the full stochastic model (solid yellow curve) and numerical calculations of the mean-field model (blue dot-dashed line) for the three modes.

### Modes of cell migration

To gain a global view of the migration behavior, we first present the phase diagram of cell migration, in terms of the migration modes, as a function of the actin polymerization speed 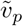, the substrate stiffness 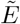, and the friction coefficient 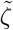 (Figure 2A). Our analysis reveals two distinct stable migration modes: gripping (fast-moving) and slipping (slow-moving), in agreement with previous experiments [38, 39]. Our model further predicts an oscillatory stick-slip mode, which has also been reported experimentally [26].

In the gripping mode, most clutches are engaged: *P*_*b*_ ≈ 1, leading to a vanishing retrograde flow and a migration speed close to 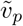 in the steady states (Figure 2D). Conversely, in the slipping mode, only a tiny fraction of clutches is engaged, resulting in significant sliding of F-actin and a low migration speed (Figure 2E), corresponding to the “frictional slippage” regime in the agent-based simulations of Ref. [26]. In the limit of low substrate friction 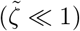, we prove that *P*_*b*_ has a lower bound in the gripping mode and is of order one. In contrast, *P*_*b*_ has an upper bound in the slipping mode that is approximately 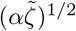 (Supplemental Material Section B).

In the stick-slip mode (e.g., the red-curve region in Figure 2C), the cell migrates in a stick-slip behavior: clutches gradually bind to the F-actin, followed by a collective rupture, producing a serrated traction force (Figure 2F). The area of the stick-slip mode in the parameter space given a fixed substrate stiffness increases as the substrate stiffness decreases (see slices of the phase diagram in Figure 2A at different 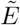 in Figure S5). We compare direct simulations of the full stochastic model (Figure 1) and numerical calculations of the mean-field model [Eqs. (1, 2)] and find they agree well, validating our mean-field assumptions (Figure 2D-F). We include the details of stochastic simulations and numerical calculations in Supplemental Material Section A.

In the following section, we show that the stick-slip mode can be further subdivided into gripping-like and slipping-like submodes. Moreover, besides the coexistence of gripping and slipping modes within the bistable phase (Figure 2B), the oscillatory stick-slip mode can also coexist with either stable mode near phase boundaries, which we discuss in a later section. Therefore, an isogenic population of cells in the same environment may exhibit subpopulations of cells, such as fast and slow-migrating cells, in agreement with experiments of freely migrating keratocytes [39].

### Migration speeds of different modes

As shown in the phase diagram (Figure 2A), the gripping mode typically occurs at low polymerization speeds. Taking the limits 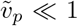 in Eq. (3) reveals that migration speed 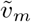 is proportional to polymerization speed 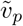 in the gripping mode:

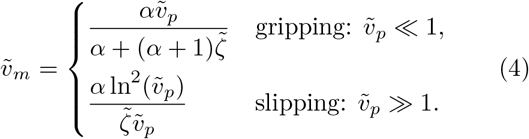

In particular, in the limit of low substrate friction 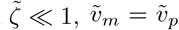: the cell migrates at its highest possible speed, independent of 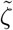, in agreement with the numerical calculations of the mean-field model (lower left panel of Figure 3B).

**FIG. 3.**
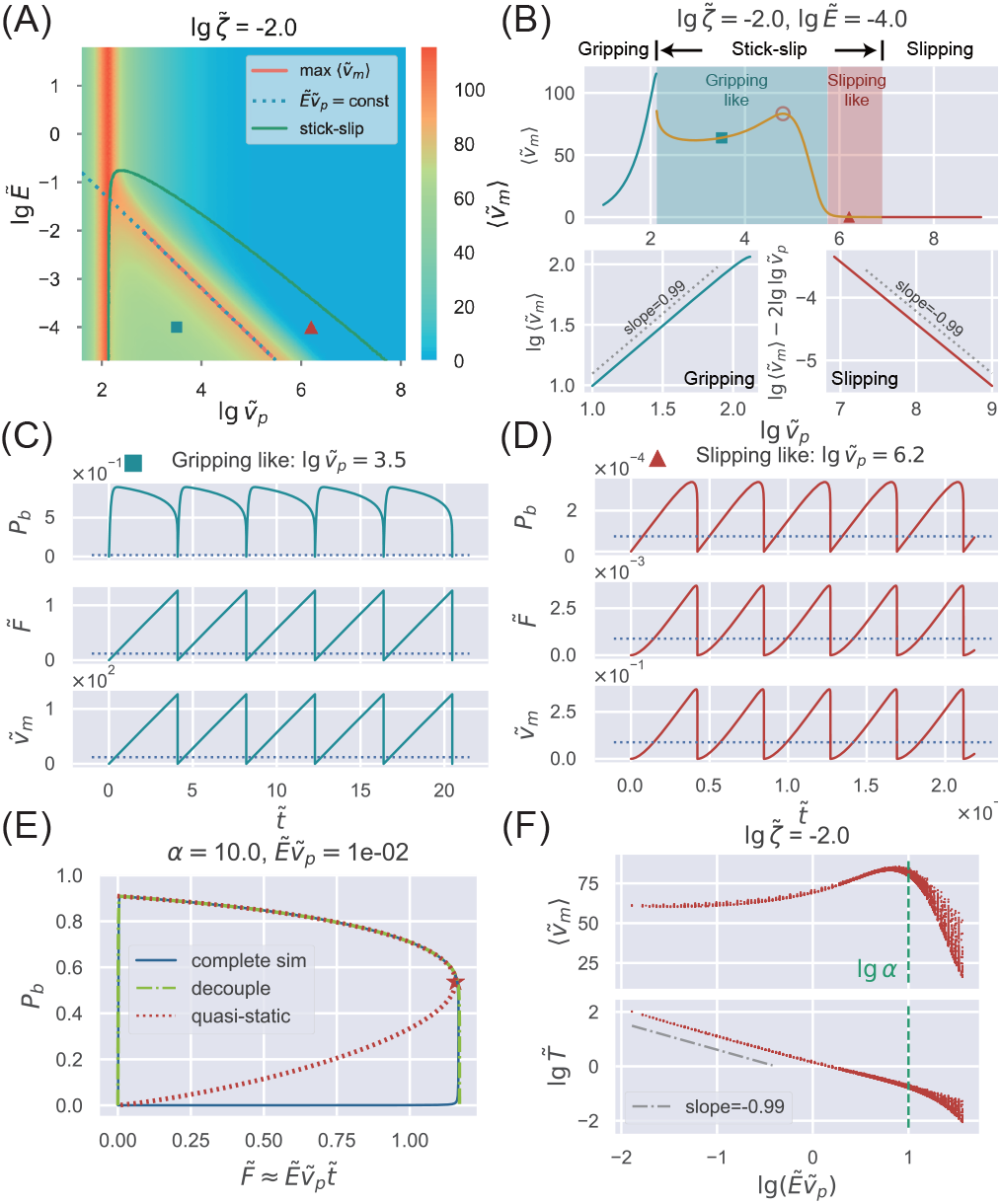
Migration speeds of different modes. (A) Heatmap of the migration speed. Averaged values during one cycle are shown for the stick-slip mode, and the green line marks the boundary of the stick-slip mode. In the stick-slip mode, the red line marks the maximum 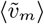. The blue dotted line is a fit to the position of maximum 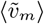 with a constant 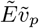. (B) A slice of the heatmap (A) along lg 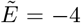. Lines of different colors denote different modes. In the stick-slip mode, the orange circle marks the maximum 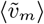. The bottom panels show the scaling relationship between 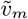 and 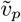 in the gripping and slipping modes, respectively. (C) A typical gripping-like stick-slip loop whose position in the parameter space is marked by the blue squares in (A) and (B). The dashed line represents the unstable fixed point of Eqs. (1, 2). (D) A typical slipping-like stick-slip loop, which corresponds to the red triangle in (A) and (B). The dashed line represents the unstable fixed point of Eqs. (1, 2) (there is only one fixed point here because 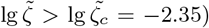. (E) The blue solid line is a direct simulation of Eqs. (1, 2). The green dot-dashed line is the same as the blue solid line but employs the decoupled dynamics approximation to Eq. (2) in the gripping-like submode. The red dotted line represents the quasi-static approximation, Eq. (7). The star marks 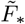 beyond which no solutions of Eq. (7) exists. (F) Parameters within the stick-slip mode of (A) are randomly sampled to examine the dependence of the stick-slip period 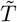 and averaged migration speed 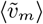 on 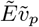. The vertical dashed line marks when 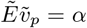. In the bottom panel, the gray dot-dashed line is a linear regression of the data for 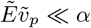.

In the slipping mode, Eq. (3) is simplified as 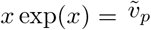 in the limit of 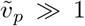, which means that 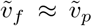 and 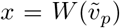 where *W* (*z*) is the root of the equation *z* = *x* exp(*x*), i.e., the Lambert W function. Therefore, 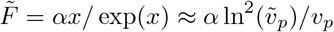, leading to the migration speed for the slipping mode in Eq. (4). Here, we use the approximation *W* (*x*) ≈ ln(*x*) for *x* ≫ 1 [48]. Surprisingly, the migration speed is inversely proportional to the actin polymerization speed 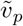 up to a logarithmic correction in the slipping mode, in agreement with numerical simulations (lower right panel of Figure 3B). The distinct behaviors of the migration speed in the 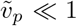 and 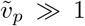 regimes suggest that the migration speed must be a non-monotonic function of the actin polymerization speed (Figure 3A, B). This prediction aligns with the biphasic relationship between traction force (proportional to migration speed in our model) and actin retrograde speed observed experimentally [49] (Figure S6).

For the stick-slip mode, we analyze the migration speed averaged over one cycle, 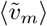. As 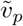 increases inside the stick-slip mode, 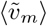 exhibits a peak, followed by a sharp but continuous decline before transitions to the slipping mode (Figure 3A, B). Intriguingly, the stick-slip mode can be divided into two distinct submodes: slipping-like and gripping-like. In the slipping-like submode, the system has negligible clutch engagement (*P*_*b*_ ≪ 1), and the system oscillates around the unstable solution of the slipping mode given by Eq. (3) (Figure 3A, D). This behavior reflects a continuous Hopf bifurcation driving the transition from the slipping mode to the slipping-like stick-slip submode (see the next section), accompanied by a continuous change in 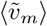 (Figure 3B).

In the gripping-like submode, corresponding to the “load-and-fail” dynamics in the agent-based simulations of Ref. [26], the system is characterized by significant clutch engagement (*P*_*b*_ ∼ 1, Figure 3C), and the ridge of maximum 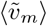 also locates in this region (Figure 3A). Interestingly, the transition from the gripping mode to the gripping-like stick-slip submode is a discontinuous Hopf bifurcation (see the next section), with a discontinuous change in 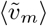 (Figure 3B).

To gain analytical insights into the stick-slip loops for the gripping-like submode, we consider the limiting case where 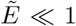 while keeping 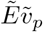 as a constant. In this case, 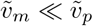 (Figure 3A), and Eq. (2) is simplified as

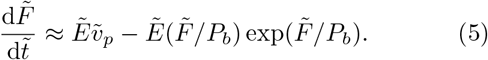

Since 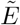 is small, the second term on the right-hand side of Eq. (5) becomes significant only when *P*_*b*_ → 0, where collective rupture occurs. Therefore, the clutch dynamics described by Eq. (5) is decoupled into two stages: a linear accumulation in the traction force 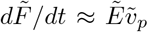, followed by a sudden rupture in all clutches when the total force reaches a threshold 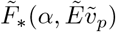, in agreement with numerical calculations of the mean-field model (Figure 3C). Given this decoupled dynamics, the average migration speed and the period of the stick-slip loop become

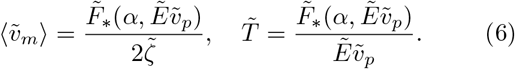

We remark that 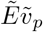 appears as an entirety in Eq. (6), which explains why the maximum 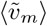 sits on the ridge of a constant 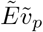 (Figure 3A).

To further verify the dependence on the product 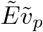, we randomly sample the values of 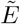 and 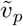 within the gripping-like stick-slip submode. We confirm that 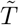 and 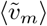, are similar as long as the product 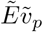 is the same (Figure 3F). Such dependence breaks down quickly as the accumulation rate of the traction force, which is 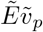, exceeds the changing rate of the binding probability, which is *α*, leading to the maximal 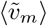 when 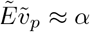. In particular, in the limit 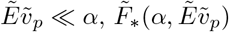 is independent of 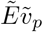; therefore, 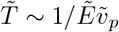 and 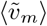 is a constant.

We highlight several intriguing predictions after translating Eqs. (6) to physical units. First, the maximum migration speed in the stick-slip mode scales as 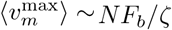 and is independent of actin polymerization speed. Second, the optimal stiffness maximizing ⟨*v*_*m*_⟩ scales as 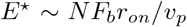 using the condition 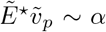 (Figure 3F). Third, the period of the stick-slip mode scales as *T* ∼ *NF*_*b*_*/*(*Ev*_*p*_). We note that some of these scaling relationships were previously identified empirically in computational studies [18, 26]. Our work provides a unifying mechanistic explanation and offers new predictions.

We seek the analytical expression of 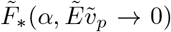. Because of the separation of the timescales that govern the dynamics of *P*_*b*_ [*α* in Eq. (1)] and the dynamics of 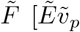 in Eq. (5)], *P*_*b*_ is always approximately at the steady-state value given the instantaneous 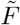:

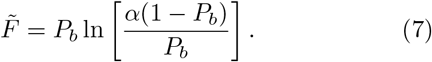

Here, *P*_*b*_ is a multi-valued function of 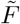. 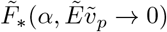 is precisely the threshold force beyond which no solutions of *P*_*b*_ exist, which is 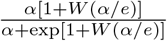 corresponding to the overall rupture in Figure 3E. To validate the quasi-static approximation, we compute the relationship between *P*_*b*_ and 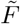 by the direct simulation of Eqs. (1, 2), a reduced simulation using Eq. (1) and the decoupled dynamics approximation 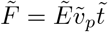, and the quasi-static results given by Eq. (7). Indeed, these three results match very well (Figure 3E).

### Boundaries between modes

The Jacobian matrix 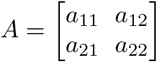 of the dynamical system [Eqs. (1, 2)] around a fixed point is

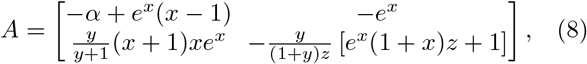

Where 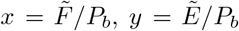, and 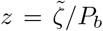 to simplify the notation. Together with the fixed point equation [Eq. (3)], we remark that, as 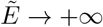, neither the fixed point location nor the Jacobian depends on the substrate stiffness 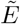. Hence, in this regime, the system dynamics is entirely independent of 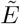, consistent with the phase diagram shown in Figure 2A. Furthermore, we find that the stick-slip mode is also absent for 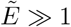 (Figure 2A).

Given *P*_*b*_ ∼ *O*(1) for the gripping mode, we find that the unstable condition for a gripping solution is *a*_11_ = −*α* + *e*^*x*^(*x* 1) *>* 0 in the limit of weak substrate stiffness 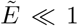 (Supplemental Material Section C), which can be written as *x >* 1 + *W* (*α/e*), where *W* (*z*) is the Lambert W function. To translate the unstable condition determined by the average force per clutch *x* to the actin polymerization speed 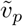, we consider two asymptotic regimes of 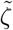. In the limit 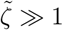, Eq. (3) is reduced to 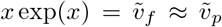; in the limit 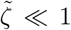, Eq. (3) is reduced to 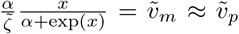. Thus, we obtain the boundary of the gripping mode in the limit 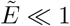 (Supplemental Material Section C):

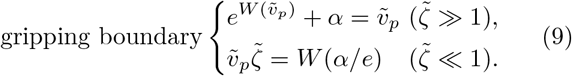

The gripping boundary at low substrate friction 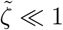, i.e., the right boundary of the bistable phase in Figure 4B, is induced by a saddle-node bifurcation. Therefore, we can get the same boundary by requiring the existence of a gripping solution in Eq. (3) without requiring the condition *E* ≪ 1 (Supplemental Material Section B). The boundary condition in physical units, *v*_*p*_*ζ/NF*_*b*_ ∼ *O*(1), has a clear physical meaning: for gripping migration, the cell must apply a traction force close to *v*_*p*_*ζ* to overcome the substrate friction since *v*_*m*_ *≈ v*_*p*_ according to Eq. (4); nevertheless, the maximum traction force has to be smaller than *NF*_*b*_, which is the maximum force all the clutches can hold.

**FIG. 4.**
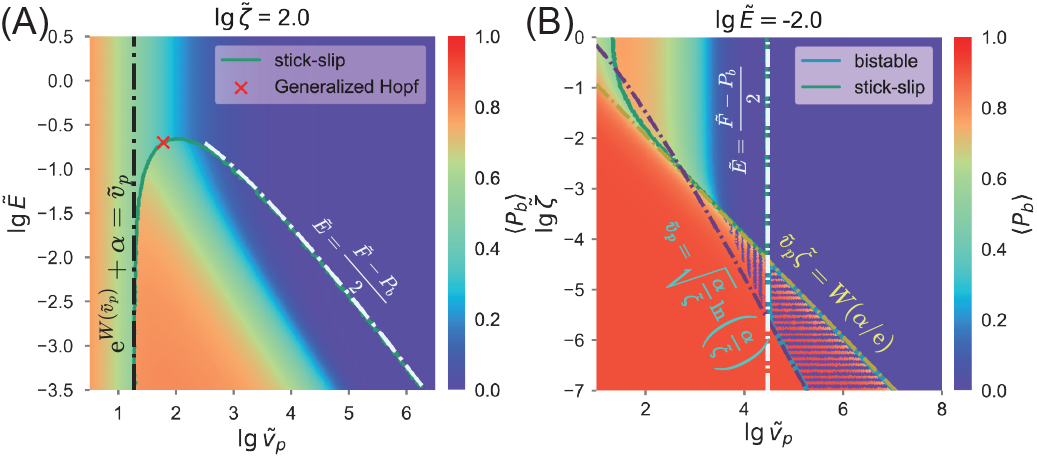
Boundaries between different migration modes. (A) Heatmap of the binding probability *P*_*b*_ at a fixed 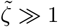. The average value over one cycle is shown for the stick-slip mode, whose boundary is marked by the green line. The black dot-dashed line represents the predicted gripping boundary [Eq. (9)]. The white line represents the predicted slipping boundary [Eq. (10)]. The red cross marks the generalized Hopf bifurcation point at which discontinuous Hopf bifurcation transitions to a continuous one. The same heatmap at a small friction coefficient *ζ < ζ*_*c*_ is shown in Figure S7. (B) The same as (A) but at a fixed 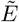. The green and blue solid lines represent the boundaries of the stick-slip mode and the bistable phase. Within the bistable phase, horizontal stripes indicate ⟨*P*_*b*_⟩ values for both stable modes. In the coexistence region of slipping-like stick-slip mode and gripping mode, ⟨*P*_*b*_⟩ is displayed as vertical stripes. The yellow dot-dashed line represents the predicted gripping boundary [Eq. (9)]. The yellow dashed line marks the saddle-node bifurcation of the slipping mode, while the vertical white line corresponds to the Hopf bifurcation, where the slipping mode becomes unstable. A heatmap as a function of 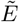 and 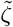 is shown in Figure S8.

Since the stability of the gripping mode is dominated by *a*_11_ in the limit of 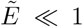, fixing 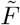 and applying a small perturbation *δP*_*b*_ is most likely to trigger instability, suggesting that the destabilization mechanism of gripping mode resembles the rupture observed in the loops of the gripping-like stick-slip submode. When the steady-state traction force 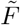 exceeds a threshold value 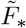, no solution of *P*_*b*_ exists. Indeed, employing 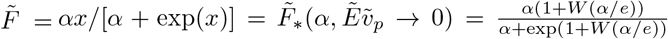 from the quasi-static approximation of the gripping-like stick-slip submode, we recover the boundary criterion for the gripping mode: *x* = 1 + *W* (*α/e*).

The slipping mode has two boundaries. The first is the boundary where stable slipping migration turns into slipping-like stick-slip, corresponding to the white dot-dashed line in Figure 4A and the vertical white dot-dashed line near lg 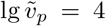 in Figure 4B. This boundary marks the instability of the slipping mode. Since clutches are barely engaged *P*_*b*_ ≪ 1 for slipping solution, we have 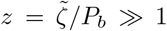. In this scenario, *a*_22_ ≈ − *y*(1 + *x*)*e*^*x*^*/*(1 + *y*). We prove that the unstable condition of the slipping mode must be governed by a Hopf bifurcation (Supplemental Material Section C and Figure S2A); therefore, the boundary of the slipping mode is tr(*A*) = 0, i.e., *y* + 1 = (*x* + 1)*/*(2 + *αe*^−*x*^), which is simplified as *y* = (*x* − 1)*/*2 using *αe*^−*x*^ *≪* 1 (recall that *P*_*b*_ = *α/*(*α* + *e*^*x*^) 1). Consequently, the unstable condition becomes 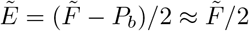, or *F/E ∼ F*_*b*_*/k*_*c*_ in physical units, which has a clear physical meaning: a significant substrate deformation relative to the maximal clutch elongation leads to stick-slip motion.

Another boundary of the slipping mode appears at the bistable phase at low substrate friction 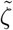, corresponding to the purple dot-dashed line shown in Figure 4B. This boundary can be derived by requiring the existence of a slipping solution in Eq. (3) when 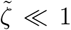, which means a saddle-node bifurcation. Specifically, it corresponds to the minimal 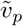 that guarantees three fixed points in Eq. (3) (Supplemental Material Section B). In summary, the boundary of the slipping mode is defined by the onset of either the instability condition or the existence condition, depending on which one is encountered first as 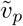 decreases from a large value:

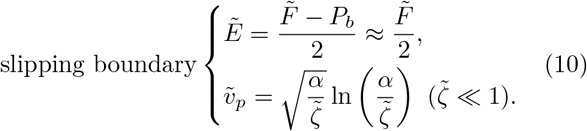

We summarize Eqs. (9, 10) in Figure 4, and leave a comprehensive discussion on the transition mechanisms between modes in Supplemental Material Section C and Table S2. We remark that at high substrate friction (Figure 4A), the entire boundary of the stick-slip mode is governed by Hopf bifurcation (Figure S2A), on which both discontinuous and continuous Hopf bifurcations occur, separated by a generalized Hopf bifurcation point (the red cross in Figure 4A). A discontinuous Hopf bifurcation induces the transition between the gripping mode and the stick-slip mode: crossing the boundary between the gripping and stick-slip modes—for instance, by increasing 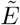—transforms the unstable gripping solution into a stable fixed point and generates an unstable limit cycle nearby (Figure S3A). The global limit cycle, corresponding to the macro stick-slip loop in the gripping-like submode, persists until it collides with the unstable limit cycle, resulting in a hysteresis loop in the migration modes (Figure S3A). Thus, even when the system exhibits only one fixed point for 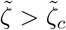, the gripping mode and the stick-slip model can still coexist near the phase boundary.

When the slipping mode becomes unstable, it undergoes a continuous Hopf bifurcation, producing a stable micro limit cycle corresponding to the slipping-like submode (Figure S3B and S4B). If a stable gripping mode also exists, the two modes coexist, as illustrated by the vertically striped region in Figure 4B. However, the limit cycle generated by the continuous Hopf bifurcation does not persist indefinitely. As 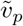 gradually decreases, it eventually collides with the middle solution of Eq. (3)—a saddle point—leading to a homoclinic bifurcation (Figure S4B), after which only the stable gripping mode remains. Consequently, this coexistence region does not extend to the saddle-node boundary of the slipping mode when 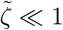 [Eq. (10)]. We remark that the slipping-like submode cannot persist if the saddle point is close to the slipping solution of Eq. (3) or if 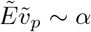, where the micro stick-slip loop gradually expands to the macro loop (see the transition between the slipping-like and gripping-like submodes in Figure 3B and Figure S4A).

## Discussion

By generalizing the motor-clutch model to the whole-cell level and using a mean-field approximation, we systematically investigate how actin polymerization speed, substrate stiffness, and friction govern cell migration. Compared with previous mean-field descriptions of the motor-clutch model [17, 24, 25, 31, 32], we take account of the dissipation of the traction force during the clutch dissociation process, i.e., the 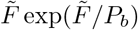 term in Eq. (2). Despite its minimal nature, our model provides key insights into the physical mechanisms of distinct migration modes, substrate stiffness sensing, and cell-to-cell variability during migration.

We identify three distinct migration modes (Figure 2A). In the gripping mode, strong clutch engagement ensures efficient traction force transmission, and the migration speed increases monotonically with the actin polymerization speed [Eq. (4)]. Conversely, the slipping mode converts actin polymerization speed primarily into retrograde flow rather than cell migration, and the migration speed decreases as the actin polymerization speed increases [Eq. (4)]. Notably, these two stable migration modes can coexist at low substrate friction but transition continuously at high friction. Our model also predicts a biphasic relationship between the migration speed and the actin retrograde speed, in agreement with previous experiments [49].

Remarkably, the migration speed in the two stable modes is independent of substrate stiffness, which means that the cell can only sense substrate stiffness in the stick-slip mode. Importantly, the stick-slip mode can be classified into two submodes: the gripping-like sub-mode, characterized by macro stick-slip loops with substantial clutch engagement and fast cell migration, and the slipping-like submode, involving small loops with negligible clutch engagement and slow migration. In between, an optimal substrate stiffness maximizes the migration speed and the traction force in the stick-slip mode. Therefore, on a substrate with a stiffness gradient, cells preferentially migrate toward this optimal stiffness, which serves as the critical threshold separating durotaxis from negative durotaxis [50]. Interestingly, the optimal stiffness scales as *E*^⋆^*v*_*p*_ ∼ *NF*_*b*_*r*_*on*_, with the maximal migration speed 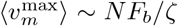, independent of the actin polymerization speed. When the substrate stiffness is significantly lower than the optimal value, the timescale of clutch binding and dissociation is much faster than that of traction force accumulation, making the gripping-like stick-slip loop a quasi-static process. In this case, the maximum traction force triggering overall rupture equals the maximal traction force achievable in the gripping mode (*F*_*_), leading to a constant average migration speed (Figure 3F).

Finally, our model predicts that an isogenic population of cells can exhibit multiple migration modes in the same mechanical environment. First, the stable gripping and slipping modes can coexist in the bistable region under low substrate friction (see the horizontal-stripe region in Figure 4B). Moreover, since the slipping mode is destabilized through a continuous Hopf bifurcation, the slipping-like stick-slip submode can coexist with the gripping mode (see the vertical-stripe region in Figure 4B). In contrast, the transition from the gripping mode to the gripping-like stick-slip submode must follow a discontinuous Hopf bifurcation under high substrate friction, leading to a hysteresis loop such that the gripping-like sub-mode can coexist with the gripping mode (Figure S3A). It will be exciting to incorporate more complex biological effects into the cell migration model in future work, such as focal adhesion reinforcement [27, 34], biochemical feedback loops on cytoskeletal activity—including Rac- and Rho-GTPase signaling [51, 52], the roles of membrane tension [21], contractility-driven cell motility [53], and periodic force dipoles in driving spontaneous polarization [54].

The research was funded by the National Key Research and Development Program of China (2024YFA0919600), National Natural Science Foundation of China (Grant No. 12474190) and Peking-Tsinghua Center for Life Sciences grants.

