## Supplementary Material for "Substrate properties and actin polymerization speed dictate universal modes of cell migration: gripping, slipping, and stick-slip"



(Dated: June 20, 2025)


### A. DETAILED DERIVATION ON THE MODEL EQUATIONS

A major distinction of our approach, compared with previous models [1–5], lies in the incorporation of traction force dissipation during the clutch dissociation process. In this section, we first formulate our stochastic framework and then derive our mean-field equations from the full stochastic model. We also present our choice for parameter normalization and estimation on typical biological systems. Finally, we outline the numerical methods for performing stochastic and ODE simulations in our work.

The system state is denoted by $\mathbf{s} = (\cdots, s_i, \cdots)$, where each binary variable $s_i$ indicates the status of clutch $i$: $s_i = 1$ if engaged, and $s_i = 0$ if unengaged. The reaction rate $r_i$ for clutch $i$ is given by:

$$r_i(s;\tau) = \begin{cases} r_{off,i}(F_i), & \text{if } s_i = 1, \\ r_{on,i}, & \text{if } s_i = 0. \end{cases} \tag{S1}$$

We remark that the reaction rates shown in Eq. (S1) are time-dependent because engaged clutches are elongated with the retrograde flow. We set the force transmitted by unengaged clutches as $F_i = 0$. Consequently, the time evolution of $F_i$ is described by:

$$\dot{F}_i = \begin{cases} k_c(v_f - \dot{x}_s), & \text{if } s_i = 1, \\ 0, & \text{if } s_i = 0, \end{cases} \tag{S2}$$

where $x_s$ denotes the substrate displacement and $v_f$ is the retrograde velocity of F-actin.

For stochastic simulations, we simulate the stochastic process described above using the Gillespie algorithm with time-dependent reaction rates [6]. Specifically, the reaction rates depend not only on the current system state $\mathbf{s}$ but also on the elapsed time $\tau$ since the last reaction event (whether binding or unbinding event) due to the retrograde flow. We define the binding fraction as $P_b = \sum_i s_i/N$ and the total traction force as $F = \sum_i F_i$. Since the elongation rates of all engaged clutches are the same, $\dot{F} = NP_b\dot{F}_i = E\dot{x}_s$ in the absence of reaction events, where $E$ is the substrate stiffness. Using $v_f = v_p - F/\zeta$ where $\zeta$ is the substrate friction coefficient, we find that $\dot{F} = NP_bk_c\left(v_p - \frac{F}{\zeta} - \frac{\dot{F}}{E}\right)$. In the absence of any reaction events, the time evolution of the total traction force becomes

$$F(\tau) = F(0) + (v_p\zeta - F(0))\left[1 - \exp\left(-\frac{NP_bk_cE}{E + NP_bk_c}\frac{\tau}{\zeta}\right)\right], \tag{S3}$$

from which we obtain the time evolution of the traction force on every engaged clutch:

$$F_i(\tau) = F_i(0) + \frac{v_p\zeta - F(0)}{NP_b}\left[1 - \exp\left(-\frac{NP_bk_cE}{E + NP_bk_c}\frac{\tau}{\zeta}\right)\right]. \tag{S4}$$

To develop a mean-field description, we focus on the time derivative of the coarse-grained state variable $P_b$: $\dot{P}_b = r_{on}(1 - P_b) - \langle r_{off,i}\rangle P_b$. Here, the angle bracket, $\langle\cdot\rangle$, indicates averaging over all engaged clutches. When neglecting the force fluctuation, $\langle r_{off,i}\rangle = r^0_{off}\exp\left(\langle F_i\rangle/F_b\right)$ leads to the expression in $\dot{P}_b$ shown in the main text. To obtain the time evolution of total traction force, $\dot{F}$, we assume that newly engaged clutches bear zero initial force. In contrast, the traction force released by unengaged clutches is approximated by the average force per engaged clutch $\langle F_i\rangle = F/(NP_b)$. Based on this, the total traction force evolves according to:

$$\Delta F = NP_bk_c(v_f - \dot{x}_s)\Delta t - NP_br_{off}\frac{F}{NP_b}, \tag{S5}$$

leading to the equation $\dot{F} = NP_bk_cv_f - \frac{NP_bk_c}{E}\dot{F} - r_{off}F$ shown in the main text.

By specifying $\bar{t} = 1/r_{off}^0$ as the time unit, $\bar{F} = NF_b$ as the force unit, and $\bar{E} = Nk_c$ as the stiffness unit, the normalization of all other physical quantities can be obtained by combining the dimensions of the above three units. We remark that these normalization factors only depends on the microscopic properties of the cells, e.g., the number of molecular clutches $N = 100$, the off-rate constant for clutch dissociation $r_{off}^0 = 0.1\ \mathrm{s}^{-1}$ [7], the characteristic clutch rupture force $F_b = 2$ pN [8] and the clutch spring constant $k_c = 5$ pN/nm [9].

| **Normalization factor** | **Reference Value** | **Dimensionless Variable** |
|---|---|---|
| Force: $\bar{F} = NF_b$ | 200 pN | $\tilde{F} = F/\bar{F}$ |
| Speed: $\bar{v} = \bar{u}/\bar{t} = F_b r_{off}^0/k_c$ | 0.04 nm/s | $\tilde{v} = v/\bar{v}$ |
| Time: $\bar{t} = 1/r_{off}^0$ | 10 s | $\tilde{t} = t/\bar{t}$ |
| Elasticity: $\bar{E} = Nk_c$ | 500 pN/nm | $\tilde{E} = E/\bar{E}$ |
| Friction: $\bar{\zeta} = \bar{F}/\bar{v} = Nk_c/r_{off}^0$ | 5000 pN·s/nm | $\tilde{\zeta} = \zeta/\bar{\zeta}$ |
| Displacement: $\bar{u} = \bar{F}/\bar{E} = F_b/k_c$ | 0.4 nm | $\tilde{x}_c = x_c/\bar{u}$ |

TABLE S1. Normalization factors and reference values for non-dimensionalization.

The complete set of equations governing the system after applying the mean-field approximation is given by:

$$\frac{\mathrm{d}P_b}{\mathrm{d}\tilde{t}} = \alpha(1 - P_b) - P_b \exp\left(\frac{\tilde{F}}{P_b}\right), \tag{S6}$$

$$(1 + P_b/\tilde{E})\frac{\mathrm{d}\tilde{F}}{\mathrm{d}\tilde{t}} = P_b(\tilde{v}_p - \tilde{F}/\tilde{\zeta}) - \tilde{F} \exp\left(\frac{\tilde{F}}{P_b}\right), \tag{S7}$$

where the system dynamics are fully governed by four independent dimensionless parameters: $\alpha = r_{on}/r_{off}^0$, $\tilde{v}_p = v_p k_c/(r_{off}^0 F_b)$, $\tilde{E} = E/(Nk_c)$, and $\tilde{\zeta} = \zeta r_{off}^0/(Nk_c)$. Among them, $\alpha$ depends solely on the microscopic properties of the ligand-receptor bond and is treated as a constant throughout our analysis. In contrast, we set $\tilde{v}_p$, $\tilde{E}$ and $\tilde{\zeta}$ as changeable parameters because they not only reflect microscopic properties but also capture distinct macroscopic influences: $\tilde{v}_p$ depends on the cytoplasmic actin polymerization speed, whereas $\tilde{E}$ and $\tilde{\zeta}$ are determined by substrate's properties.

We remark that the above system of ordinary differential equations is numerically stiff, meaning it involves both rapidly varying components (e.g., the sudden force drops during overall rupture events in the stick-slip mode) and slowly varying components (e.g., the gradual traction force accumulation in gripping-like stick-slip submode). This numerical stiffness poses challenges for numerical integration using standard explicit methods, which would require prohibitively small time steps to maintain stability and accuracy. To address this, we employ numerical solvers with automatic stiffness detection and adaptive method switching. Specifically, we utilize the Rodas4P algorithm, as implemented in Ref. [10], and the LSODA algorithm, as described in Ref. [11].

### B. PROPERTIES OF STABLE MIGRATION MODES

A stable migration mode is characterized by a constant migration speed, a constant binding probability $P_b$, and a constant traction force $\tilde{F}$. By setting the time derivatives in Eqs. (S6, S7) to zero, we obtain the following fixed point equation, which is Eq. (3) in the main text:

$$\tilde{v}_p = \frac{\alpha}{\tilde{\zeta}} \frac{x}{\alpha + \exp(x)} + x \exp(x), \tag{S8}$$

where we define $x = \tilde{F}/P_b$ for notational simplicity.

We first demonstrate that the fixed point equation Eq. (S8) possesses at most one inflection point. Taking the second derivative of the right-hand side yields the condition for inflection points:

$$\frac{\tilde{\zeta}}{\alpha}(e^x + \alpha)^3 + \left(e^x \frac{x-2}{x+2} - \alpha\right) = 0. \tag{S9}$$

Differentiating Eq. (S9) once more with respect to $x$ reveals that the left-hand side is a monotonically increasing function. Hence, Eq. (S8) can have at most one inflection point, two extrema, and thus up to three fixed points. As shown in Figure S1A, the necessary and sufficient condition for the emergence of an inflection point is $\tilde{\zeta} < \tilde{\zeta}_c$, under which the right-hand-side of Eq. (S8) is non-monotonic.

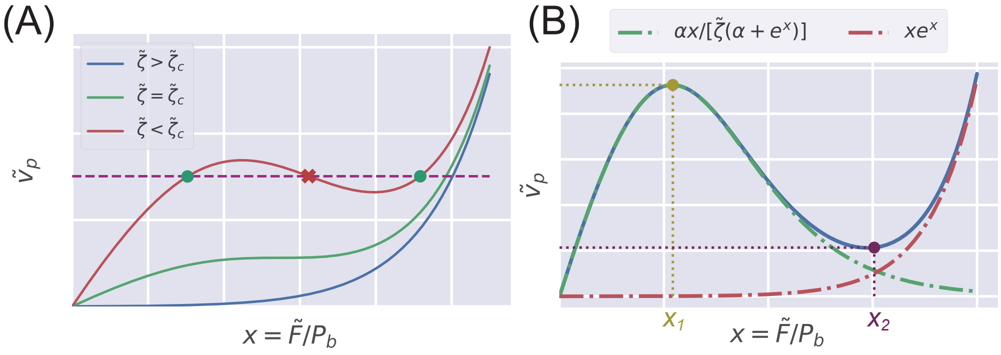


FIG. S1. **Plot of the fixed point equation Eq. (S8).** (A) When the substrate friction $\tilde{\zeta}$ falls below a critical threshold $\tilde{\zeta}_c$, and $\tilde{v}_p$ lies within an appropriate range, Eq. (S8) may admit three roots. The green dots indicate the stable fixed points, while the red cross marks the unstable fixed point. (B) In the limit $\tilde{\zeta} \ll 1$, the two contributions to Eq. (S8) are shown separately. The green dot-dashed line represents the first term on the right-hand side (corresponding to $\tilde{v}_m$), and the red dot-dashed line represents the second term (corresponding to $\tilde{v}_f$). The yellow and purple dots denote analytical estimates of the first and second extrema, $x_1$ and $x_2$, respectively, given by Eqs. (S11) and (S14).

We prove that if the system exhibits three roots (i.e., $\tilde{\zeta} < \tilde{\zeta}_c$), the middle solution must be unstable, regardless of the value of $\tilde{E}$. By treating $x = \tilde{F}/P_b$ as the independent variable, the normalized dynamics described by Eq. (S7) can be rewritten as:

$$\frac{\mathrm{d}x}{\mathrm{d}\tilde{t}} = \frac{1}{1 + P_b/\tilde{E}} \left( \tilde{v}_p - \frac{P_b x}{\tilde{\zeta}} - x \exp(x) \right) - x \left[ \alpha \left( \frac{1}{P_b} - 1 \right) - \exp(x) \right]. \tag{S10}$$

For a migration mode to be stable, perturbations in any direction of $(\delta P_b, \delta x)$ must decay over time. In particular, when the system is perturbed along the direction where $\mathrm{d}P_b/\,\mathrm{d}\tilde{t} = 0$, the stability along the $x$ direction is governed entirely by the first term on the right-hand side of Eq. (S10). Stability is maintained only if the term $P_b x/\tilde{\zeta} + x \exp(x)$ is locally increasing with respect to $x$. As shown in Figure S1A, the middle solution is unstable because the right-hand side of Eq. (S8) decreases as $x$ increases.

### Asymptotic behaviors in the $\tilde{\zeta} \ll 1$ limit

We study the properties of the two stable migration modes in the limit $\tilde{\zeta} \ll 1$. As established in the preceding discussion, the gripping solution lies to the left of the first extremum of Eq. (S8), and the slipping solution to the right of the second extremum. Both solutions fall within regions where $\tilde{v}_p$ increases locally with respect to $x$ (Figure S1A). To characterize the first extremum denoted by $x_1$, we approximate:

$$\frac{\alpha}{\tilde{\zeta}} \frac{x_1}{\alpha + \exp(x_1)} \approx \tilde{v}_p, \quad (x_1 - 1)\exp(x_1) = \alpha. \tag{S11}$$

The second equation follows from setting the derivative of the right-hand side of Eq. (S8) to zero. Solving Eq. (S11) yields $x_1 = 1 + W(\alpha/e)$, where $W(z)$ is the Lambert W function, the inverse function of $xe^x$. Therefore, for a valid gripping solution, it must hold that

$$\tilde{v}_p \tilde{\zeta} \leq \frac{\alpha x_1}{\alpha + \exp(x_1)} x_1 - 1 = W(\alpha/e). \tag{S12}$$

As the left-hand side of the first equation in Eq. (S11) is indeed $\tilde{F}/\tilde{\zeta}$, the extremum also maximizes the traction force. Consequently, in the gripping mode, the migration speed $\tilde{v}_m$ increases monotonically with $\tilde{v}_p$ up to the point of maximal traction. Additionally, we can estimate the lower bound for the binding probability as:

$$P_b \geq \frac{\alpha}{\alpha + \exp(x_1)} = P_{b*} \sim O(1). \tag{S13}$$

For the second extremum $x_2$, we estimate:

$$\frac{\alpha}{\tilde{\zeta}} \frac{x_2}{\exp(x_2)} + x_2 \exp(x_2) \approx \tilde{v}_p, \quad \frac{\alpha}{\tilde{\zeta}} \approx \exp(2x_2)\frac{x_2+1}{x_2-1} \tag{S14}$$

Similarly, the second expression in Eq. (S14) results from setting the derivative of the right-hand side to zero. Here, we assume $\exp(x_2) \gg \alpha$, which will be verified subsequently. We define $x_{2,0} = \frac{1}{2}\ln(\alpha/\tilde{\zeta})$ as the leading-order approximation of the second expression in Eq. (S14), and Taylor expand $x_2$ around $x_{2,0}$ to obtain the first-order correction:

$$\delta x_2 = -\frac{x_{2,0}-1}{x_{2,0}^2 - 2}, \quad \delta \tilde{v}_p = 2\sqrt{\frac{\alpha}{\tilde{\zeta}}}\delta x_2 \tag{S15}$$

Since $\delta\tilde{v}_p/\tilde{v}_p = \delta x_2/x_{2,0} \sim O(x_{2,0}^{-2})$, ignoring higher order corrections and approximating $x_2$ as $x_{2,0}$ can well estimate the second extremum of Eq. (S8) (Figure S1B). This leads to the condition for the existence of a slipping mode:

$$\tilde{v}_p \geq \sqrt{\frac{\alpha}{\tilde{\zeta}}} \ln\left(\frac{\alpha}{\tilde{\zeta}}\right). \tag{S16}$$

Finally, as $\tilde{\zeta} \to 0$, we have $\exp(x_{2,0}) = \sqrt{\alpha/\tilde{\zeta}} \gg \alpha$, thus confirming the validity of the earlier assumption $\exp(x_2) \gg \alpha$. The condition for the slipping migration mode can be written as $\tilde{v}_f = x\exp(x) > \frac{\alpha x}{\tilde{\zeta}\exp(x)} = \tilde{v}_m$, which has a clear physical interpretation: $\tilde{v}_p$ is not primarily converted into $\tilde{v}_m$, but rather dissipated by the unbinding process, that is, the cell is slipping. Accordingly, an upper bound for the binding probability in the slipping mode can be estimated as

$$P_b = \frac{\alpha}{\alpha + \exp(x)} < \sqrt{\alpha\tilde{\zeta}}, \tag{S17}$$

given the fact that $\exp(2x) > \alpha/\tilde{\zeta}$. This result shows that most clutches are unengaged in the slipping mode.

### C. BOUNDARIES BETWEEN MIGRATION MODES

In this section, we investigate the asymptotic behaviors of the migration mode boundaries as a function of the three parameters $\tilde{E}$, $\tilde{\zeta}$, and $\tilde{v}_p$ as they approach zero or $+\infty$. Table S2 provides a summary of the boundaries between different migration modes. We begin with the Jacobian of the system defined by Eqs. (S6, S7), expressed as

$$A = \begin{bmatrix} a_{11} & a_{12} \\ a_{21} & a_{22} \end{bmatrix} = \begin{bmatrix} -\alpha + e^x(x-1) & -e^x \\ \frac{y}{y+1}(x+1)xe^x & -\frac{y}{(y+1)z}\left[e^x(1+x)z + 1\right] \end{bmatrix}, \tag{S18}$$

where we define $x = \tilde{F}/P_b$, $y = \tilde{E}/P_b$, and $z = \tilde{\zeta}/P_b$ for short of notation. Since the linear stability analysis focuses on perturbations around stable migration modes, Eq. (S8) has already been substituted into Eq. (S18) to simplify the expression.

#### Asymptotic behaviors at $\tilde{E} \ll 1$

In this subsection, we focus on the boundary of the gripping mode in the limit of $\tilde{E} \ll 1$. By definition, in the gripping mode, $P_b \sim O(1)$; therefore, the Jacobian matrix in Eq. (S18) becomes

$$A = \begin{bmatrix} a_{11} & a_{12} \\ \tilde{E}a'_{21} & \tilde{E}a'_{22} \end{bmatrix} = \begin{bmatrix} -\alpha + e^x(x-1) & -e^x \\ \frac{\tilde{E}}{P_b}(x+1)xe^x & -\frac{\tilde{E}}{P_b}e^x(1+x) - \frac{\tilde{E}}{\tilde{\zeta}} \end{bmatrix}. \tag{S19}$$

| Condition | Criteria | Bifurcation | Mode | Expression | Reference |
|---|---|---|---|---|---|
| $\tilde{\zeta} \ll 1$ | Extreme points of Eq. (S8) | Saddle-node | Slipping | $\tilde{v}_p = \sqrt{\alpha/\tilde{\zeta}} \ln\left(\alpha/\tilde{\zeta}\right)$ | Eq. (S16) |
| | | | Gripping | $\tilde{v}_p \tilde{\zeta} = W(\alpha/e)$ | Eq. (S12) |
| $\tilde{E} \ll 1$ | Perturbation along $(\delta P_b, 0)$ <br> $a_{11} = 0$ | Hopf | Gripping | $\tilde{v}_p \tilde{\zeta} = W(\alpha/e),\ (\tilde{\zeta} \ll 1)$ | Eq. (S24) |
| | | | | $e^{W(\tilde{v}_p)} + \alpha = \tilde{v}_p,\ (\tilde{\zeta} \gg 1)$ | Eq. (S23) |
| $\tilde{v}_p \gg 1$ | Eigenvalues of $A$ <br> $\mathrm{tr}\,(A) = 0$ | Hopf | Slipping | $\tilde{E} = \frac{1}{2}(\tilde{F} - P_b)$ | Eq. (S31) |
| $\tilde{\zeta} \gg 1$ | | | Gripping | $e^{W(\tilde{v}_p)} + \alpha = \tilde{v}_p,\ (\tilde{E} \ll 1)$ | Eq. (S30) |

TABLE S2. A summary of boundary conditions discussed in our work.

Since $\tilde{E}$ is small, the eigenvalues can be expanded as:

$$\lambda_1 = a_{11} + \frac{a_{12}a'_{21}}{a_{11}}\tilde{E} + O(\tilde{E}^2), \tag{S20a}$$

$$\lambda_2 = \left(a'_{22} - \frac{a_{12}a'_{21}}{a_{11}}\right)\tilde{E} + O(\tilde{E}^2). \tag{S20b}$$

As our analysis focuses on the gripping boundary, $a_{12}a'_{21} \sim O(1) < 0$ and $a'_{22} \sim O(1) < 0$ always hold, while $a_{11}$ may change the sign according to the value of $x$ at the fixed point. The stability of the gripping solution requires both eigenvalues $\lambda_1$ and $\lambda_2$ to be negative. From the eigenvalue expansion [Eq. (S20)], this condition reduces to $a_{11} < 0$, yielding the inequality

$$(x - 1)\exp(x) < \alpha. \tag{S21}$$

Importantly, this inequality is not only necessary but also sufficient for generating a gripping boundary. For a fixed but arbitrary small $a_{11} < 0$, the sign of $a_{11}$ also ensures $\lambda_2$ to be negative.

As the stability of the gripping mode fully depends on the sign of $a_{11}$, the system is most susceptible to perturbations in the direction of $[\delta P_b,\ 0 \cdot \delta\tilde{F}]$. Therefore, the instability of the gripping mode is equivalent to require the equation

$$\frac{dP_b}{d\tilde{t}} = \alpha(1 - P_b) - P_b \exp\left(\frac{\tilde{F}_{\text{const}}}{P_b}\right) \tag{S22}$$

has no stable fixed point under a constant traction force $\tilde{F}_{\text{const}}$. Since the right-hand side of Eq. (S22) is negative as $P_b \to 0$ and $P_b \to 1$, the fixed points of Eq. (S22) always appear in pairs, one stable and one unstable. Although this phenomenon resembles a saddle-node bifurcation, where a stable node and an unstable saddle are simultaneously created or annihilated as a system parameter passes through a critical value, we later show that the boundary at $\tilde{E} \ll 1$ is actually a Hopf bifurcation, and resolves this apparent contradiction at the end of this subsection.

Thus, the instability condition for the gripping mode can be interpreted as follows: when the applied load $\tilde{F}_{\text{const}}$ exceeds a critical threshold $\tilde{F}_*$, Eq. (S22) admits no steady-state solution; that is, the unbinding rate surpasses the binding rate at all times. This physical scenario aligns with the overall rupture behavior observed in gripping-like stick-slip mode discussed in the main text. Consequently, both the threshold forces for the overall rupture in the stick-slip loop and a stable gripping migration correspond to the same critical value of $x = 1 + W(\alpha/e)$.

To derive a concise expression for the phase boundary, we consider the limiting cases $\tilde{\zeta} \to +\infty$ and $\tilde{\zeta} \to 0$. These limits differ in the asymptotic behavior of $\tilde{v}_p$ as a function of $x$ in Eq. (S8).

1. In the limit $\tilde{\zeta} \to +\infty$, Eq. (S8) simplifies to $x\exp(x) = \tilde{v}_f \approx \tilde{v}_p$. Thus, the boundary condition becomes

$$e^{W(\tilde{v}_p)} + \alpha = \tilde{v}_p, \tag{S23}$$

which defines a surface $\tilde{v}_p(\alpha)$ orthogonal to the $\tilde{v}_p$ axis (see the black dot-dashed line in Figure 4A of the main text).

2. In the limit $\tilde{\zeta} \to 0$, Eq. (S8) reduces to $\tilde{v}_m \approx \tilde{v}_p$. In this case, the boundary is approximated by:

$$\tilde{v}_p \tilde{\zeta} \approx \tilde{F}_* = W(\alpha/e), \tag{S24}$$

which is characterized by a boundary with a slope of -1 in the $\lg \tilde{\zeta}$—$\lg \tilde{v}_p$ plane (see the yellow dot-dashed line in Figure 4B of the main text.)

Finally, we explain why the boundary of the gripping mode consists of Hopf bifurcation points in the limit $\tilde{E} \ll 1$. A subtle point in the derivation of Eq. (S20) is that when the eigenvalue is expanded as a Taylor series in $\tilde{E}$, the expansion becomes invalid as $a_{11} \to 0$, since the radius of convergence vanishes. Specifically, for a function of the form $\sqrt{1 + k\epsilon}$, the radius of convergence in $\epsilon$ is $1/|k|$. Thus, the expansion in Eq. (S20) is valid only when:

$$\tilde{E} < \left| \frac{a_{11}^2}{4a_{12}a'_{21} - 2a_{11}a'_{22}} \right| \sim O(a_{11}^2). \tag{S25}$$

Importantly, Eq. (S25) specifies not only the convergence radius of the eigenvalue expansion but also the condition for the discriminant of the characteristic polynomial of the Jacobian matrix $A$ to be positive in the limit $\tilde{E} \ll 1$.

In Figure S2A, B, we show the eigenvalue properties as a function of $\tilde{v}_p$ and $\tilde{E}$ for the cases $\tilde{\zeta} \gg 1$ and $\tilde{\zeta} \ll 1$, respectively. In both regimes, as long as $\tilde{E}$ is sufficiently small, as the actin polymerization speed $\tilde{v}_p$ increases, the eigenvalues always transition from two real values to a pair of complex conjugates. Moreover, the region in which the eigenvalues are complex conjugates narrows as $\tilde{E}$ decreases. Consequently, for a fixed small $\tilde{E}$, there always exists a sufficiently small $|a_{11}|$ such that the expansion in Eq. (S20) breaks down, and the eigenvalues become a pair of complex conjugates—indicating a Hopf bifurcation at the gripping boundary.

To conclude, a precise interpretation of the results in this subsection is as follows: for a given $a_{11} > 0$, there always exists a sufficiently small $\tilde{E}$ that renders the gripping solution unstable; conversely, for a given $a_{11} < 0$, there always exists a sufficiently small $\tilde{E}$ that stabilizes the gripping solution. Therefore, we can use the condition $a_{11} = 0$ to define the gripping phase boundary in the limit $\tilde{E} \to 0$.

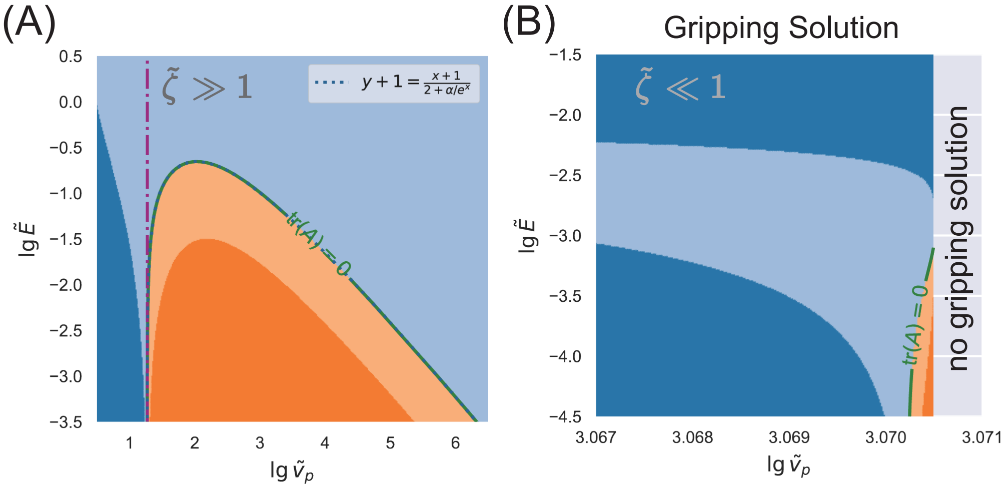


FIG. S2. **Eigenvalue properties of the Jacobian matrix** $A$. In this figure, both eigenvalues given by Eq. (S18) have negative real parts in the blue region, indicating local stability. In contrast, at least one eigenvalue has a positive real part in the orange region, indicating local instability. Furthermore, dark-colored areas represent real eigenvalues, while light-colored areas represent complex conjugate pairs. (A) The eigenvalue properties in the limit $\tilde{\zeta} \gg 1$. The dotted curve shows the theoretical prediction from Eq. (S27), which aligns well with the boundary given by $\mathrm{tr}\,(A) = 0$. In this panel, $\lg \tilde{\zeta} = 8$. (B) The eigenvalue properties in the limit $\tilde{\zeta} \ll 1$. In this regime, Eq. (S8) yields three roots, and we focus on the Jacobian at the gripping solution. We expand the abscissa to better illustrate the eigenvalue properties near the boundary. The blank region indicates the absence of a gripping solution. In this panel, $\lg \tilde{\zeta} = -3$. One should note that an extremely thin dark-blue region exists between the light-blue region and the blank region, which is invisible in this plot.

### Asymptotic behaviors at $z \to \infty$

In this subsection, we focus on the scenario where $z \to +\infty$ in Eq. (S18). Under this condition, the fixed-point equation [Eq. (S8)] reduces to $\tilde{v}_p \approx x\exp(x)$, and the Jacobian matrix elements simplify to:

$$a_{21} \approx \frac{y(1+x)}{1+y}\tilde{v}_p, \quad a_{22} \approx -\frac{y(1+x)}{1+y}e^x \tag{S26}$$

We first assume that, at the boundary of the stick-slip mode, the Jacobian eigenvalues form a complex conjugate pair, $\lambda_{1,2} = a \pm b\mathrm{i}$. Therefore, for the corresponding migration mode to be stable, the trace of matrix $A$ must satisfy

$$\operatorname{tr} A \approx -\alpha + (x-1)e^x - \frac{y}{y+1}e^x(1+x) < 0 \quad \Rightarrow \quad y+1 = \frac{x+1}{2+\alpha/e^x} \tag{S27}$$

Here, the second expression of Eq. (S27) corresponds to the boundary condition of the stick-slip mode. This transition, where a pair of complex conjugate eigenvalues crosses the imaginary axis, corresponds to a Hopf bifurcation. To confirm that the eigenvalues indeed become complex at the boundary, we evaluate the discriminant of the characteristic polynomial of the Jacobian:

$$\Delta = (\lambda_1+\lambda_2)^2 - 4\lambda_1\lambda_2 = -4\det A \approx -4\frac{(1+x)y}{(1+y)^2}(x-y)e^{2x} < 0, \tag{S28}$$

where the inequality follows from the boundary condition:

$$y+1 = \frac{x+1}{2+\alpha/e^x} \quad \Rightarrow \quad x = (2+\alpha/e^x)(y+1) - 1 > 2y+1. \tag{S29}$$

Numerical calculations confirm that Eq. (S27) accurately captures the phase boundary in the entire range of $\lg\tilde{v}_p$ in the limit of $z \to +\infty$ (Figure S2A).

We next study the asymptotic behaviors in the two simplified cases corresponding to the gripping and slipping boundaries, which provide further analytical insight. Notably, along the boundary between the gripping mode and the stick-slip mode, since $P_b \sim O(1)$, $z \gg 1$ implies $\tilde{\zeta} \gg 1$. In contrast, for the boundary between the slipping mode and the stick-slip mode, since $P_b \ll 1$, $z \gg 1$ can be satisfied without requiring $\tilde{\zeta} \gg 1$.

1. As shown in Figure. S2A (also see Figure 4A in the main text), the left boundary between the gripping mode and the stick-slip mode asymptotically approaches a vertical line, becoming nearly parallel to the $\lg\tilde{E}$-axis. This suggests an appropriate asymptotic limit for analyzing this boundary is $\tilde{E} \ll 1$. After setting $y \to 0$, Eq. (S27) simplifies to

$$(x-1)\exp(x) = \alpha, \tag{S30}$$

   which is indeed the same as the boundary condition we derive in the previous subsection in the $\tilde{\zeta} \gg 1$ limit [Eq. (S23)]. This boundary condition suggests that the binding probability remains finite, with $P_b \sim O(1)$ as $\tilde{E} \to 0$, thereby justifying our previous choice of the limit $y = \tilde{E}/P_b \to 0$, as it indeed captures the onset of the gripping mode. Additionally, this result indicates that in the asymptotic limits $\tilde{E} \to 0$ and $\tilde{\zeta} \to +\infty$, the binding probability of the gripping mode approaches the same lower bound as in the opposite limit $\tilde{\zeta} \to 0$ [Eq. (S13)].

2. For the slipping boundary, we consider the limit $\tilde{v}_p \to +\infty$, in which $P_b \sim W(\tilde{v}_p)/\tilde{v}_p \to 0$, and the assumption $z \to +\infty$ is automatically satisfied. Thus, the expression we derive below is valid both in the large $\tilde{v}_p$ and large $\tilde{\zeta}$ limits. At this boundary, we obtain $y = \frac{1}{2}(x-1)$, or equivalently:

$$\tilde{E} = \frac{1}{2}(\tilde{F} - P_b) \approx \frac{\tilde{F}}{2}, \tag{S31}$$

   where the approximation holds because $\tilde{F}/P_b = x \approx \ln(\tilde{v}_p) \gg 1$.

#### Asymptotic behaviors at $\tilde{\zeta} \ll 1$

In the discussion of steady-state properties in the previous section, we show that when $\tilde{\zeta} \to 0$, the fixed point equation has three roots. For the gripping solution, the binding fraction $P_b$ has a lower bound $P_b \geq P_{b*}$. Therefore, taking the limit $\tilde{\zeta} \to 0$ is equivalent to setting $z \to 0$ in Eq. (S18). In this case, only $a_{22}$ in the Jacobian matrix diverges,

$$a_{22} \approx -\frac{y}{z(1+y)}e^x \sim O(z^{-1}), \tag{S32}$$

while the other elements are all $\sim O(1)$. The eigenvalues of the Jacobian at this time are:

$$\lambda_1 = a_{22} + O(z^1), \tag{S33a}$$

$$\lambda_2 = a_{11} + O(z^1). \tag{S33b}$$

Since $a_{22} < 0$, $\lambda_1$ is always a negative real number. Therefore, the stability of the gripping solution is also determined by $a_{11}$. Interestingly, $a_{11} = 0$ corresponds exactly to the condition of extreme point in the fixed point equation Eq. (S11). The eigenvector for $\lambda_2$ is parallel to the $[\delta P_b, 0 \cdot \delta\tilde{F}]$ direction, corresponding to the direction of least instability.

It is important to note that the expansion in Eq. (S33) does not suffer from the convergence radius issue encountered in Eq. (S25). Indeed, for fixed $\tilde{E}$ (even if very small), sufficiently small $\tilde{\zeta}$ ensures that the Jacobian matrix possesses two real eigenvalues when $a_{11} = 0$. This gives rise to a new paradox: in the joint limit $\tilde{E} \ll 1$ and $\tilde{\zeta} \ll 1$, does the gripping boundary lose stability via a Hopf bifurcation or a saddle-node bifurcation? As shown in Figure S2B, both transition mechanisms coexist, and the outcome depends on the order in which the limits are taken.

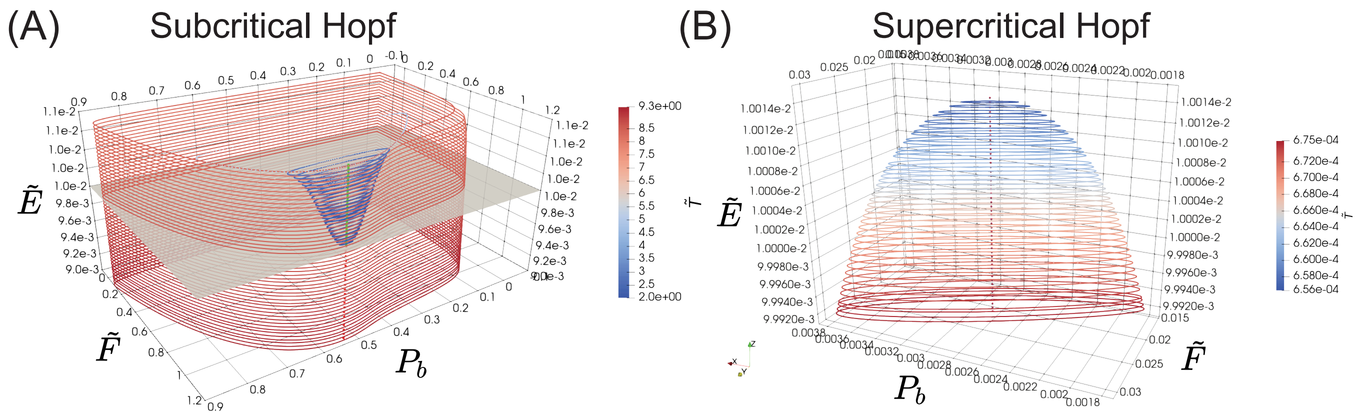


FIG. S3. **Limit cycles arising from Hopf bifurcations of different stable migration modes.** (A) With $(\alpha, \tilde{\zeta}, \tilde{v}_p) = (10, 100, 19.4)$ fixed, varying only the substrate stiffness $\tilde{E}$ (z-axis) does not affect the fixed point $(P_b, \tilde{F})$ in the xy-plane. For $\tilde{E} > 0.01$, the gripping solution is stable (green solid line), while for $\tilde{E} < 0.01$, it becomes unstable (red dotted line). The closed orbits parallel to the $xy$-plane represent limit cycles at different $\tilde{E}$ values, with color indicating their oscillation periods. A subcritical (i.e., discontinuous) Hopf bifurcation gives rise to an unstable local limit cycle near the stable fixed point. The global limit cycle (associated with gripping-like stick-slip) persists beyond the bifurcation point, resulting in a coexistence region of stable gripping and stick-slip modes, as well as a hysteresis loop in migration modes. (B) For the supercritical (i.e., continuous) Hopf bifurcation from the slipping solution, we fix $(\alpha, \tilde{\zeta}, \tilde{v}_p) = (10, 0.01, 29200)$. When $\tilde{E} < 1.0015 \times 10^{-2}$, the slipping solution becomes unstable (red dotted line), and a stable limit cycle with increasing amplitude emerges, which means a continuous transition from the slipping mode to the slipping-like submode.

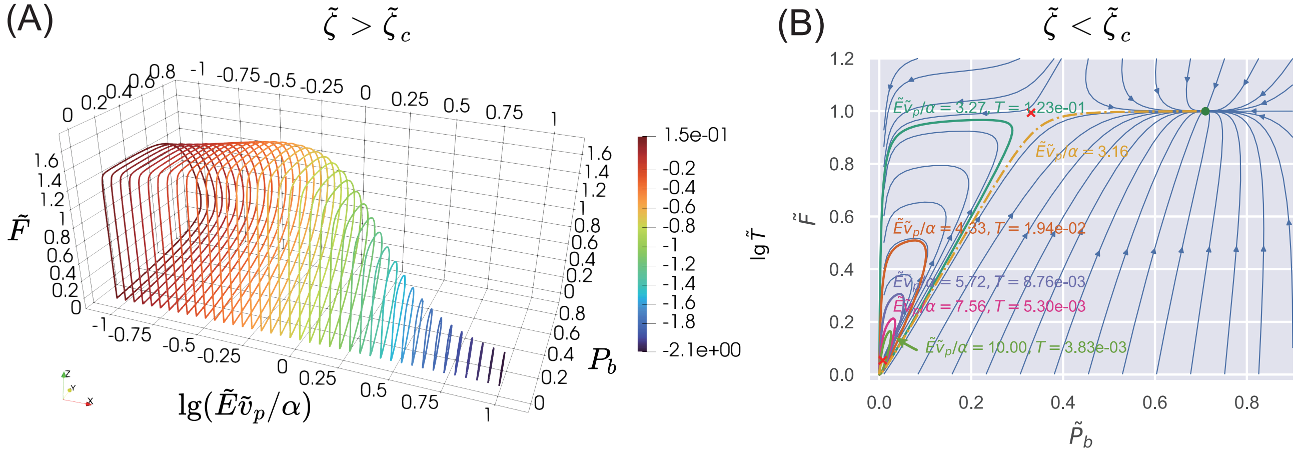


FIG. S4. **Gradual transition from a slipping-like to a gripping-like stick-slip loop.** (A) With $(\lg \alpha, \lg \tilde{\zeta}, \lg \tilde{v}_p) = (1, -2, 3.5)$ fixed, we vary the substrate stiffness $\tilde{E}$ and focus on the evolution of the stick-slip limit loop. Since $\tilde{\zeta} > \tilde{\zeta}_c$, Eq. (S8) yields a single, unstable fixed point enclosed by a stable limit cycle. Orbit colors indicate the oscillation period $\log \tilde{T}$. As $\tilde{E}\tilde{v}_p \sim \alpha$, the slipping-like stick-slip loop gradually transforms into a gripping-like one, accompanied by a significant increase in the oscillation period. (B) For $(\lg \alpha, \lg \tilde{\zeta}, \lg \tilde{v}_p) = (1, -4, 4)$, a similar expansion of the slipping-like stick-slip loop is observed as $\tilde{E}$ increases (closed orbits marked with different colors). In this case, $\tilde{\zeta} < \tilde{\zeta}_c$ and Eq. (S8) yields three fixed points, with the gripping solution being the only stable one. The middle solution (red cross near $(P_b, \tilde{F}) = (0.35, 1)$) corresponds to a saddle point, which can be seen from the blue streamplot. As the limit cycle expands, it eventually collides with the saddle point, leading to a homoclinic bifurcation. Beyond this point, the limit cycle vanishes, and trajectories near the slipping solution are attracted to the stable gripping state (e.g., the yellow dot-dashed line corresponding to $\tilde{E}\tilde{v}_p/\alpha = 3.16$).

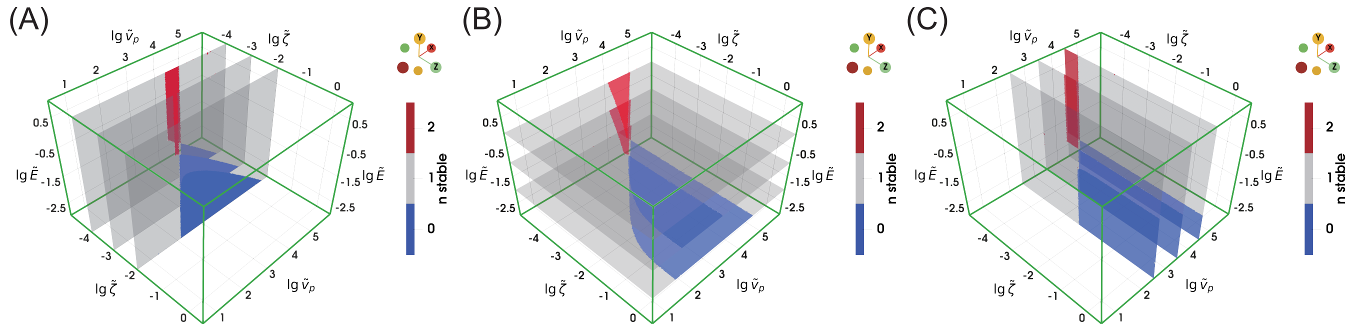


FIG. S5. **Slices of the phase diagram in three parameter planes.** Slices of the phase diagram from Figure 2A of the main text are shown in the $\lg \tilde{E}$—$\lg \tilde{v}_p$ plane (A), $\lg \tilde{\zeta}$—$\lg \tilde{v}_p$ plane (B), and $\lg \tilde{E}$—$\lg \tilde{\zeta}$ plane (C). Colors indicate the number of stable migration modes under the corresponding parameter combinations. Notably, panel (B) shows that the stick-slip region expands as the substrate stiffness $\tilde{E}$ decreases.

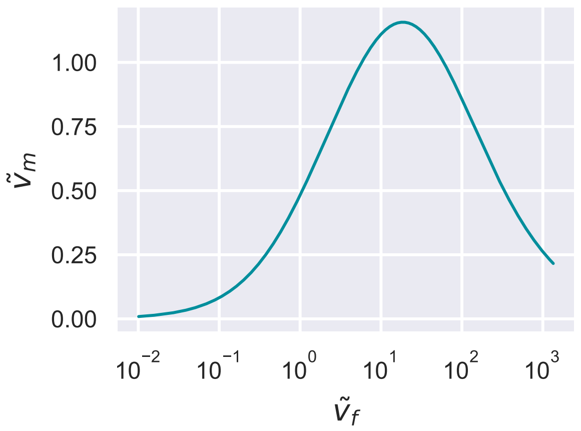


FIG. S6. **Biphasic relationship between the cell migration speed $\tilde{v}_m$ and the F-actin retrograde speed $\tilde{v}_f$.** We take $\tilde{\zeta} = 1$ and $\tilde{E} = 10^4$ such that $\tilde{\zeta} > \tilde{\zeta}_c$ and $\tilde{E}$ is sufficiently large, allowing a continuous transition from the gripping to the slipping mode without entering the stick-slip regime.

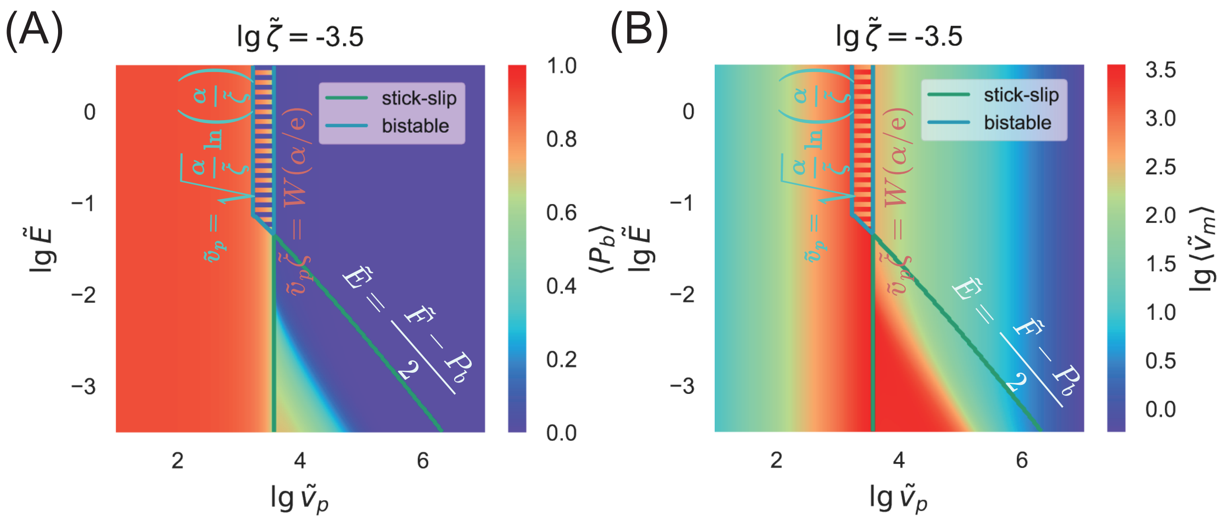


FIG. S7. **Heatmap of migration modes on** $\lg \tilde{E}$—$\lg \tilde{v}_p$ **plane.** The averaged value of $P_b$ (panel A) and $\tilde{v}_m$ (panel B) over one cycle is shown for the stick-slip mode, whose boundary is marked by a green line. The oblique boundary indicates where the slipping mode becomes unstable and transitions into a slipping-like submode, as described by Eq. (S31). In contrast to the case presented in Figure 4A of the main text, here $\lg \tilde{\zeta} = -3.5 < \lg \tilde{\zeta}_c$ is used, resulting in a bistable phase. The vertical boundary near $\lg \tilde{v}_p = 3.5$ corresponds to the existence condition of the gripping mode [Eq. (S12)], and the vertical one near $\lg \tilde{v}_p = 3$ corresponds to the slipping mode [Eq. (S16)]. To simultaneously visualize both slipping and gripping solutions within the bistable phase, we use alternating horizontal stripes to represent their corresponding values of $\langle P_b \rangle$ (panel A) and $\langle \tilde{v}_m \rangle$ (panel B). A blue line marks the boundary of the bistable phase.

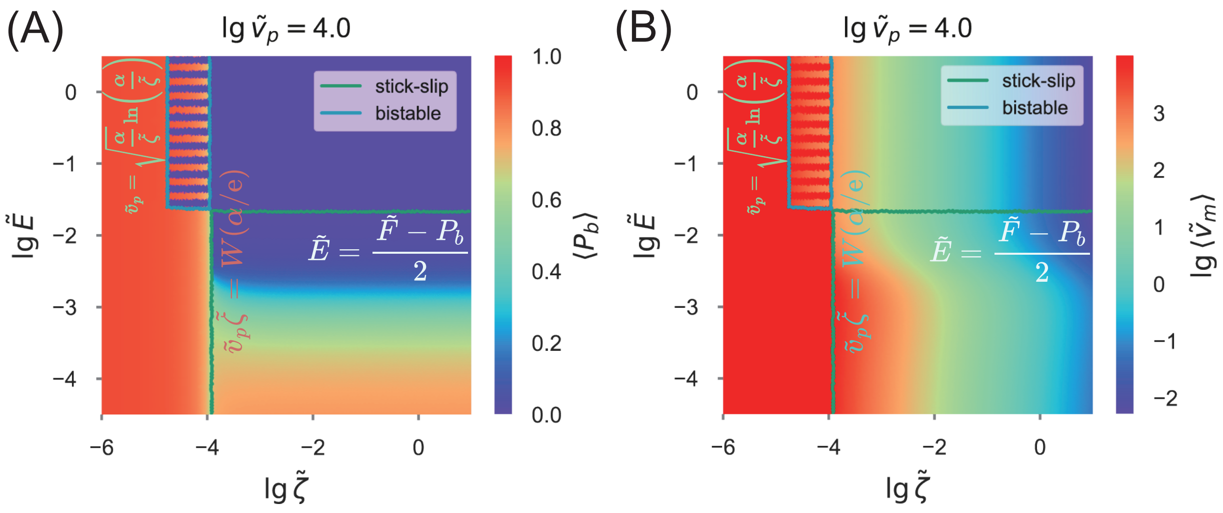


FIG. S8. **Heatmap of migration modes on** $\lg \tilde{E}$—$\lg \tilde{\zeta}$ **plane.** The averaged value of $P_b$ (panel A) and $\tilde{v}_m$ (panel B) over one cycle is shown for the stick-slip mode, whose boundary is marked by a green line. The nearly horizontal boundary near $\lg \tilde{E} = -1.7$ indicates where the slipping mode becomes unstable and transitions into a slipping-like submode, as described by Eq. (S31). The bistable phase is bounded by the blue line: the vertical boundary near $\lg \tilde{\zeta} = -5$ corresponds to the existence condition of the slipping mode [Eq. (S16)], and the vertical one near $\lg \tilde{\zeta} = -4$ corresponds to the gripping mode [Eq. (S12)]. To simultaneously visualize both slipping and gripping modes within the bistable phase at small $\tilde{\zeta}$, we use alternating horizontal stripes to represent their corresponding values of $\langle P_b \rangle$ (panel A) and $\langle \tilde{v}_m \rangle$ (panel B).
